# Aberrant cortical spine dynamics after concussive injury are reversed by integrated stress response inhibition

**DOI:** 10.1101/2022.05.31.494250

**Authors:** Elma S. Frias, Mahmood S. Hoseini, Karen Krukowski, Maria Serena Paladini, Katherine Grue, Gonzalo Ureta, Kira D.A. Rienecker, Peter Walter, Michael P. Stryker, Susanna Rosi

**Author notes:** Corresponding Authors: Susanna Rosi, Ph.D., Altos Labs, Redwood City, CA 94065; Michael Stryker, PhD, 675 Nelson Rising Lane, Room 535 Department of Physiology, University of California San Francisco San Francisco, CA 94143; Peter Walter, Ph.D., Altos Labs, Redwood City, CA 94065. **Author Contributions:** E.S.F., M.S.H., K.K., M.P.S., and S.R. designed research; E.S.F., M.S.H., M.P., G.U., and K.D.A.R. performed research; E.S.F., K.K., M.P., K.G., P.W., M.P.S., and S.R. analyzed data; and E.S.F., M.P.S., P.W., and S.R. wrote the paper. **Data Sharing:** Code used in the analysis is available at the sites noted in the text and references. Data and other documentation are available from the authors at the addresses above.

## Abstract

Traumatic brain injury (TBI) is a leading cause of long-term neurological disability in the world and the strongest environmental risk factor for the development of dementia. Even mild TBI (resulting from concussive injuries) is associated with a >2-fold increase in the risk of dementia onset. Little is known about the cellular mechanisms responsible for the progression of long lasting cognitive deficits. The integrated stress response (ISR), a phylogenetically conserved pathway involved in the cellular response to stress, is activated after TBI, axsnd inhibition of the ISR — even weeks after injury — can reverse behavioral and cognitive deficits. However, the cellular mechanisms by which ISR inhibition restores cognition are unknown. Here we used longitudinal two-photon imaging *in vivo* after concussive injury in mice to study dendritic spine dynamics in the parietal cortex, a brain region involved in working memory. Concussive injury profoundly altered spine dynamics measured up to a month after injury. Strikingly, brief pharmacological treatment with the drug-like small-molecule ISR inhibitor ISRIB entirely reversed the structural changes measured in the parietal cortex and the associated working memory deficits. Thus, both neural and cognitive consequences of concussive injury are mediated in part by activation of the ISR and can be corrected by its inhibition. These findings suggest that targeting ISR activation could serve as a promising approach for the clinical treatment of chronic cognitive deficits after TBI.

**Significance Statement:** After traumatic brain injury, temporary pharmacological inhibition of the integrated stress response (ISR), with a small-molecule inhibitor (ISRIB), rescued long lasting trauma-induced cognitive deficits. Here, we found that ISRIB treatment rapidly and persistently reversed the aberrant changes in cortical spine dynamics in the parietal cortex while rescuing working memory deficits. These data suggests that the link between the ISR and memory function involves, at least in part, changes in neuronal structure. Targeting ISR activation could serve as a promising approach for the clinical treatment of chronic cognitive deficits after brain injuries.

## Introduction

Traumatic brain injury (TBI) is defined as the disruption of normal brain function after an impact (a bump, blow, or jolt) to the head that produces direct damage from mechanical forces and indirect damage from secondary responses. Between 1.5 – 3.8 million people experience TBI each year in the United States alone (*1, 2*). At least 75% of TBI-diagnosed injuries are considered “mild” (defined as concussive injuries without loss of consciousness) (*1*). TBI is a serious health concern due to chronic behavioral and cognitive impairments that affect the quality of life of millions of individuals (*3–7*). TBI is also the strongest environmental risk factor for the development of dementia (*4, 5*). While pre-clinical experimental studies are steadily increasing, little is known about the cell-specific changes in the brain that are responsible for the persistent cognitive deficits that develop after even mild TBI. With no identified mechanisms, there are no treatments to prevent or mitigate deficits resulting from TBI.

The closed head injury (CHI) mouse model for mild TBI produces cognitive deficits comparable to those observed after concussive injuries in humans, such as working memory dysfunction (*8–10*). A type of working memory, known as short-term memory, is dependent on multiple cortical regions, including prefrontal and parietal cortex (*11–14*). All types of memory (including short-term memory) are thought to be encoded in connections among neurons at synapses (*15–17*). Most excitatory synapses connect to excitatory neurons via dendritic spines, which are specialized protuberances containing the postsynaptic receptors for neurotransmitters (*18–21*). Different types of cortical neurons have between one hundred and a few thousand dendritic spines. Memory (both formation and maintenance) depends on synaptic plasticity, i.e., the capacity of synapses to undergo lasting biochemical and morphological changes in strength in response to neuronal activity and neuromodulators. Changes in synaptic plasticity are associated with lasting changes in the structure of dendritic spines (structural plasticity) (*22–27*).

Dendritic spines consist of a spine head connected to the dendritic shaft by a spine neck (*28, 29*). When connecting stable, functional synapses, dendritic spines may be stable for months to years. Functional synapses require dendritic spines to have a bulbous spine head, as they harbor neurotransmitter receptors, intracellular signaling molecules, ribosomes that mediate local protein synthesis, and a highly active cytoskeleton meshwork (*30–32*). Spine necks are barriers that filter the electrical component of synaptic signals and amplify spine head depolarization, as well as limit the diffusion of intracellular messengers from the spine head into the dendrite (*33, 34*).

Dendritic spines have captured the attention of neuroscientists for over 100 years because they are the elements of synaptic connectivity that are visible in the light microscope. Spines are characterized by their dynamics, that is, their structural plasticity, density, and morphology. Spine structural plasticity describes the formation and elimination of spines that take place over hours or days, which is important for the development and proper function of the central nervous system (*30, 35*). Spine density is measured as the number of spines per measured length of dendrite and is used as a measure of the total synaptic strength received by a dendrite (*28*). Spine morphology describes the shape and size of dendritic spines (*36*). The regulation of each of these factors is essential for stable synaptic maintenance and transmission and thus cognitive and executive function, such as working memory (*26*).

The pioneering work from Lendvai et al, Gruzendler et al., Trachtenberg et al., and Holtmaat et al in the early 2000s demonstrated that it was possible to track the same spine in the living brain over a long period of time using two-photon imaging *in vivo* (*37–40*). From such time-lapse imaging studies, a new picture of spines began to emerge. Dendritic spines could be seen *in vivo* to change their shape and size (morphology), to form or disappear across an animal’s life span (dynamics), and to be responsive to the animals’ experience and environment. Later studies have shown that spine morphology and dynamics vary among neuronal types and across developmental stages (*19*). The balance between spines forming or disappearing determines spine density. Currently, changes in spine morphology, dynamics, and density are accepted markers for changes in synaptic strength or synaptic presence.

The integrated stress response (ISR) is a universally conserved intracellular signaling network that responds to a variable environment to maintain homeostasis (*41, 42*). Four specialized kinases (PERK (PKR-like ER kinase), PKR (double-stranded RNA-dependent protein kinase), GCN2 (general amino acid control nonderepressible), and HRI (heme-regulated inhibitor 2)) converge on the phosphorylation of a single serine on the α subunit of the translation initiation factor eIF2 (*42, 43*). PERK, PKR, and GCN2 are expressed in the mammalian brain (*43*). The phosphorylation-mediated central regulatory step leads to a reduction in global protein translation and to the translational up-regulation of a select subset of mRNAs, such as activating transcription factor 4 (ATF4) (*44, 45*). ISR-mediated reprogramming of translation maintains or reestablishes physiological homeostasis; however, chronic ISR activation can also create maladaptive cellular and functional changes (*46*). Previous work from our lab demonstrated that pharmacological inhibition of the ISR using the drug-like small molecule ISR inhibitor (ISRIB) is sufficient to reverse electrophysiological impairments, as well as chronic cognitive and behavioral deficits in different rodent TBI models (*8, 9*). The cellular mechanisms by which ISR inhibition restores neuronal function, and ultimately cognitive function, remain unknown. In the current study, we used longitudinal two-photon imaging *in vivo* to reveal cortical spine dynamics weeks after a single mild closed-head concussive injury and measured the effect of temporary ISR inhibition weeks after the injury on both spine dynamics and working memory.

## Results

### Concussive injury alters cortical spine dynamics

To determine the effect of concussive injury on spine dynamics and density, we performed longitudinal transcranial two-photon imaging *in vivo* using Thy1-YFP-H transgenic mice. In this transgenic mouse line, YFP is expressed in a small subset of layer V pyramidal neurons that extend their apical dendrites to layer I/II of the cortex (*47*). We first subjected mice to a single mild concussive closed-head injury (CHI), a reproducible and translational model for concussive injury (*48*). To perform CHI, animals were secured to a stereotaxic frame with non-traumatic ear bars.

We surgically exposed the skull and induced a single bilateral closed-head impact injury with a pneumatic piston over the parietal cortex. No animal had a fractured skull after injury. After impact, the scalp was sutured. Sham controls received the same surgery but no impact. This concussive injury produced no changes in neuronal numbers in the parietal cortex, as measured by quantification of NeuN-positive neuronal nuclei (**Supp Figure 1**). Eight days later, we implanted a head fixation device and cranial window directly above the parietal cortex (encompassing the impacted region).

To collect longitudinal data on dendritic spines, we imaged apical dendritic segments in 38 mice for 4-5 sessions over 8 to 15 days (starting at day 11 post-CHI or post-sham surgery), with the final imaging on day 25 (green arrowheads in **Figure 1**). We analyzed each dendritic segment independently and imaged between 40 and 57 segments in each of the 4-5 imaging sessions. We first quantified the number of spines per dendrite on day 11 (first imaging session) and then identified new spines present on day 13, 15, 18, and 25 (subsequent imaging sessions). To this end, we gave each spine a unique identifier (represented by X, Y, Z coordinates). Similarly, we next quantified the number of spines per dendrite on day 11 and asked whether these spines were still present on subsequent imaging sessions. Based on these measurements, we defined (i) **“spine formation”** as the fraction of new spines formed in a new location, i.e., spines in locations that were not present in the first imaging session and (ii) **“spine elimination”** as the fraction of pre-existing spines eliminated, both measurements in reference to baseline day 11 (**Figure 2A**). Representative images for these analyses are shown in **Figure 2B**.

**Figure 1.**
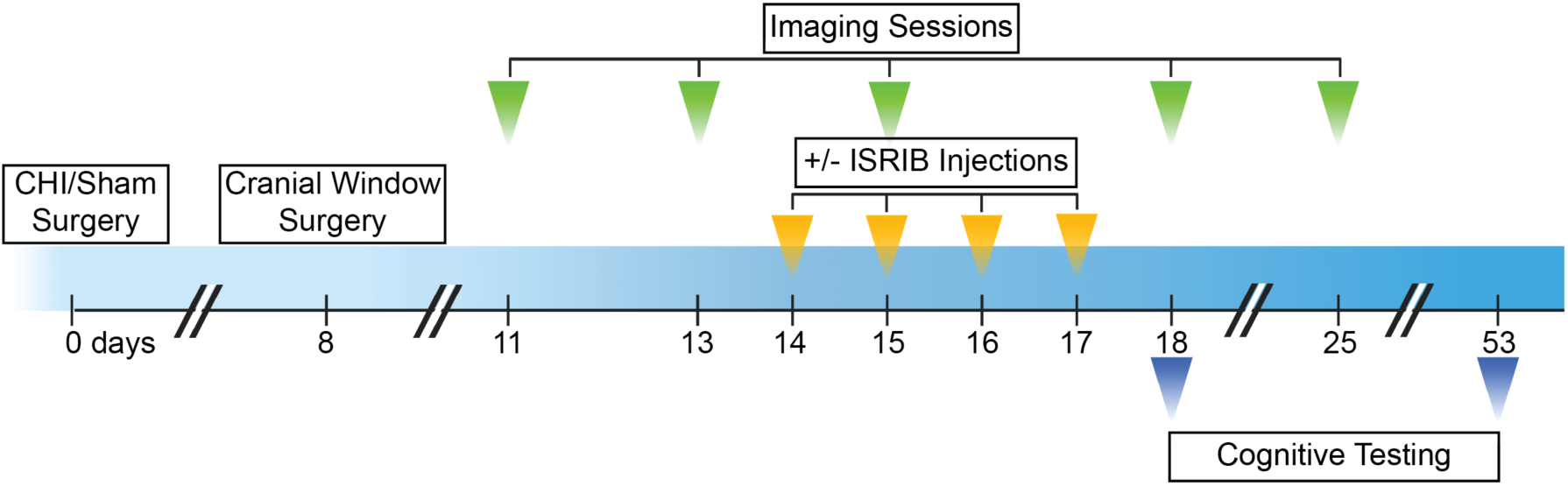
Schematic representation of experiment design. Experimental design for surgeries, imaging, ± ISRIB treatment paradigm, and cognitive testing. For the imaging experiments, adult Thy1-YFP-H transgenic male mice were subjected to CHI or sham surgery (day 0). On day 8 a cranial window was implanted directly above the parietal cortex (encompassing the impacted region). Mice were imaged for four to five sessions starting at day 11 with the final imaging day on days 18 or 25. Mice received four daily intraperitoneal injections of ISRIB or Vehicle (2.5mg/kg) starting on day 14. For the cognitive testing, adult WT male mice were first subjected to CHI or sham surgery (day 0), then received ± ISRIB treatment starting on day 14, and lastly underwent cognitive testing on day 18 and 53.

**Figure 2.**
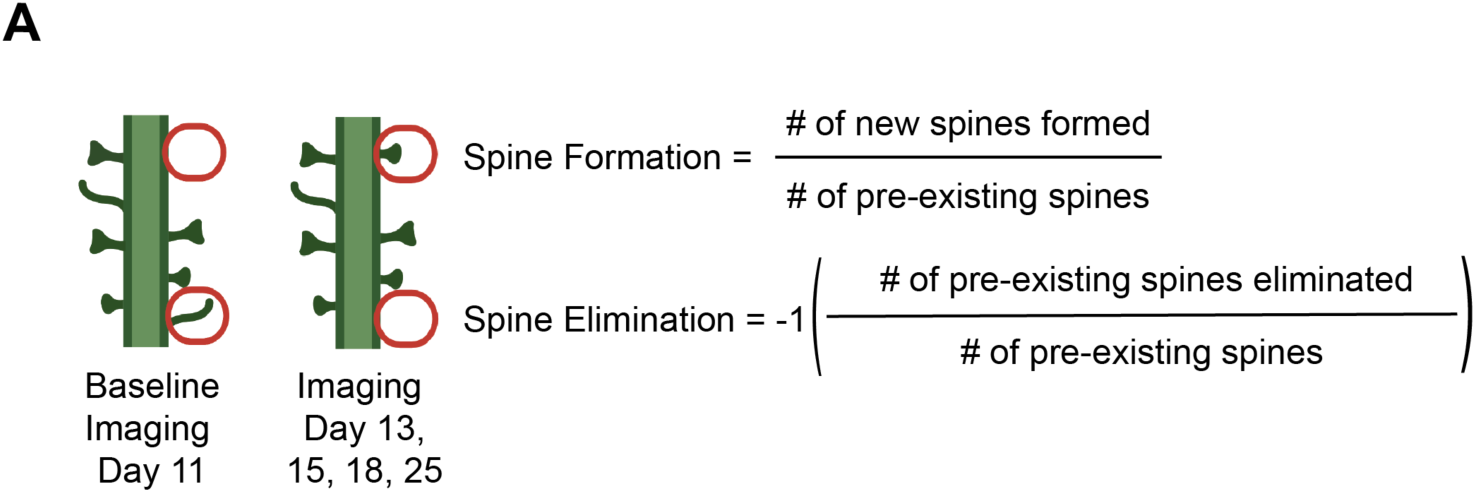
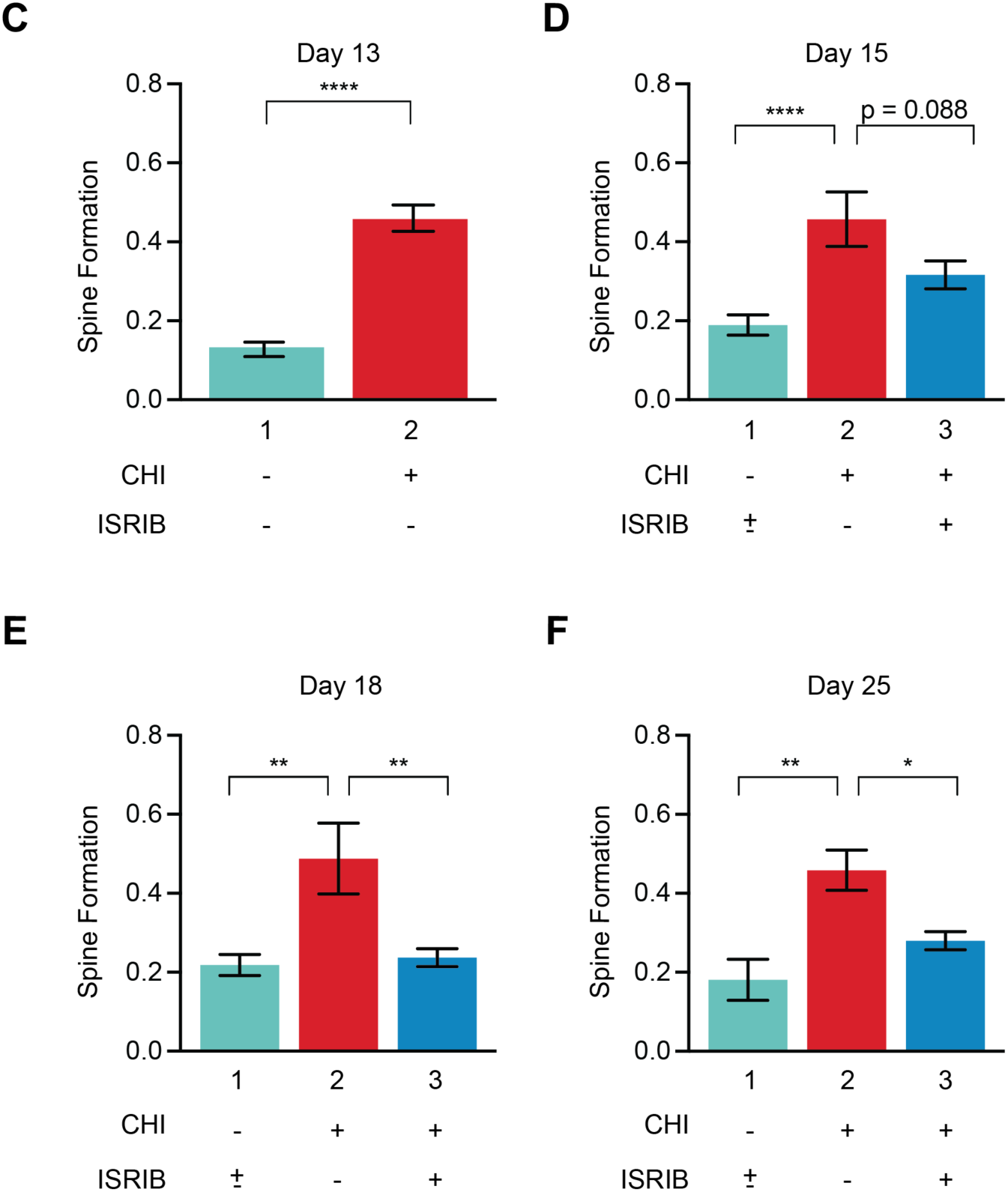
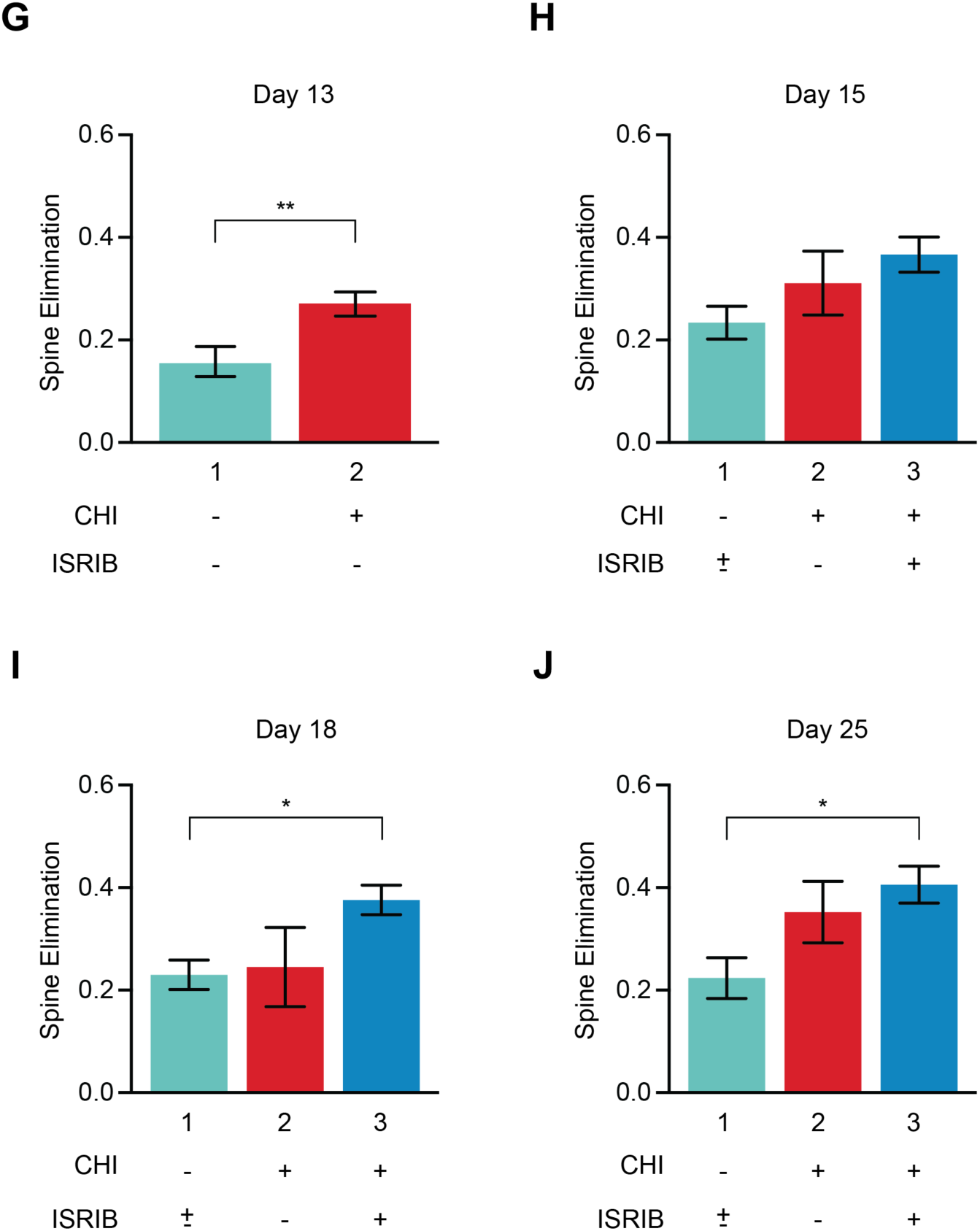
Cortical spine dynamics are altered by concussive injury and restored by ISR inhibition. **(A)** Analysis schematic depicting how spine formation and spine elimination were scored. Imaging days 13, 15, 18, and 25 were compared to baseline imaging day 11. **(B)** Representative images. White arrowheads show spines identified on one dendritic segment at baseline; blue arrowheads indicate new spines; orange arrowheads indicate spines that were eliminated. **(C-J)** Spine formation and spine elimination for ± CHI and ± ISRIB treatment. Sham ± ISRIB treatment showed no significant difference and were pooled. Statistics were calculated between the experimental groups and the pooled sham ± ISRIB treatment. **(C)** Spine formation at day 13 post- injury (dpi) for sham (Bar 1) and CHI (Bar 2) mice. The number of animals in each group was n = 13 and 23, respectively. **(D-F)** Spine formation at days 15, 18, and 25 dpi for sham ± ISRIB (Bar 1) and CHI (Bar 2), and CHI + ISRIB treatment (Bar 3). ISRIB treatment had no effect on sham control mice (Supp Figure 4). The number of animals in each group was n = 13, 12, and 11, respectively (note that the 23 animal cohort analyzed in (B) was spilt into the ± ISRIB groups). **(G- J)** same as (C-F) but spine elimination was scored. **Statistics:** Analysis was done by Unpaired T- test (C, G) or Ordinary one-way ANOVA followed by multiple comparisons using Tukey-post hoc (D-F, and H-J). p < 0.05 (*), p < 0.005 (**), p < 0.0001 (****), as indicated in the figure. All data are means ± SEM.

In a few cases (< 30%), in which a spine appeared to be eliminated but reappeared in a subsequent session in the same location, that spine was counted as if it were a reappearance of the original spine. We counted all protrusions from the dendrites greater than 0.55 µm in length as spines regardless of their morphology. In total, we analyzed a length of at least 3500 µm of dendrites (40-57 segments, each 50-100 µm (+/- 2 µm)) in length per mouse, amounting to 200- 600 spines, per experimental group. Analysis was done semi-automated and blinded to treatment and surgical condition. A detailed summary of the data sampling is listed in **Table 1**.

**Table 1.**
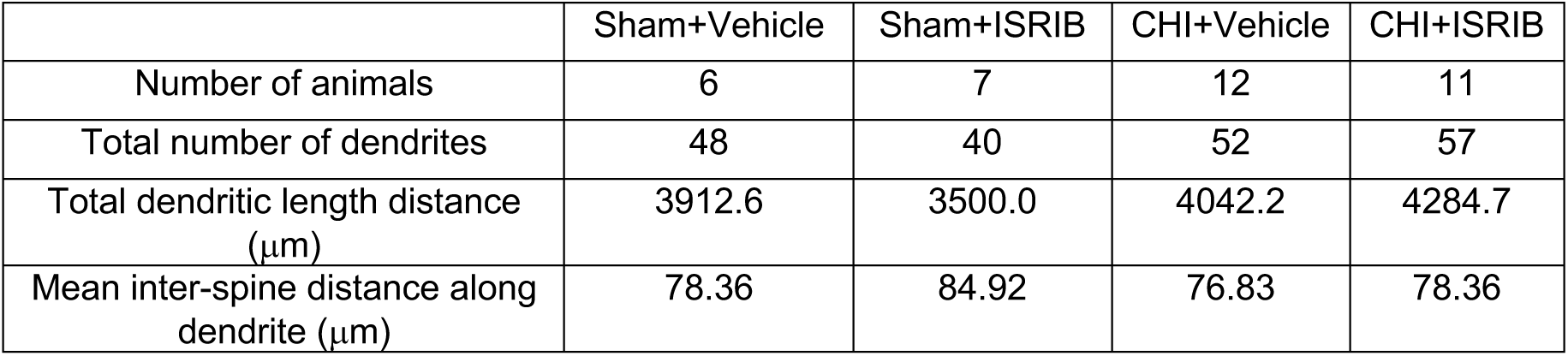
Detailed summary of the data sampling for dendritic spine analysis.

|  | Sham+Vehicle | Sham+ISRIB | CHI+Vehicle | CHI+ISRIB |
| --- | --- | --- | --- | --- |
| Number of animals | 6 | 7 | 12 | 11 |
| Total number of dendrites | 48 | 40 | 52 | 57 |
| Total dendritic length distance<br>($\mu\text{m}$) | 3912.6 | 3500.0 | 4042.2 | 4284.7 |
| Mean inter-spine distance along<br>dendrite ($\mu\text{m}$) | 78.36 | 84.92 | 76.83 | 78.36 |

At 13 days post-injury, mice that received CHI showed a significant increase in spine formation as compared to control mice that received sham surgery. This increase in spine formation was observed between each subsequent imaging sessions. As expected, control mice maintained a ratio of ∼0.2 new spines formed per pre-existing spine (*49*), which increased to ∼0.5 after injury (compare bars 1 and 2, **Figure 2C-F**). In contrast, to their significant increase in spine formation, mice that received CHI showed no significant change in spine elimination as compared to control mice (compare bars 1 and 2, **Figure 2G-J**). We further validated these conclusions and excluded that any measured changes were due to phototoxicity by reverse-time analysis, using day 18 as the baseline imaging day (**Supp Figure 2**). We conclude that concussive injury massively increases spine formation. As we show below, many of these newly formed spines are transient and are quickly eliminated, and as such do not factor into our spine elimination score, which scores changes in pre-existing spines from the baseline images.

### ISR inhibition persistently reverses the aberrant spine dynamics measured after concussive injury

We previously showed that ISR inhibition alleviates cognitive deficits after CHI (*8*). Therefore, we next tested whether pharmacological blockage of the ISR using the drug-like small-molecule ISR inhibitor ISRIB would modify the aberrant spine dynamics observed after CHI. To this end, we administered ISRIB (or vehicle) on four consecutive days starting at day 14 post-CHI or post-sham surgery (yellow arrowheads in **Figure 1**). ISRIB concentrations in the brain measured at 3 and 17 h following the last injection were consistent with earlier findings (**Supp Figure 3**) (*50*).

We observed that at day 15, only one day after the first injection, spine formation in CHI mice that received ISRIB treatment was restored toward control levels and was maintained similar to control levels thereafter (compare bars 2 and 3, **Figure 2C-F**). Importantly, we observed no effects of ISRIB treatment on spine measurements in control mice that received sham surgery (compare bars 1 and 2 for days 15-25, **Supp Figure 4**). In contrast to its effect on spine formation, ISRIB treatment only modestly affected spine elimination (**Figure 2G-J**). As above, these relationships were also observed in the reverse analysis using day 18 as the baseline imaging day (**Supp Figure 2**). Thus, ISRIB treatment reversed the CHI-induced increase in spine formation back to the levels of uninjured mice. This restorative effect persisted for at least one week after ISRIB treatment was ended on day 17 post-surgery (compare bars 2 and 3, **Figure 2F**), that is, long after the drug (T_1/2_ ∼8 hours, (*50*)) had cleared from the system.

From the longitudinal data, we also measured spine density, calculated as the number of spines per micrometer of dendritic length and observed no significant differences among groups.

### ISR inhibition normalizes the CHI-induced short-lived newborn spines

We next used the longitudinal data to examine the persistence of the new spines formed after concussive injury and whether they were long- or short-lived. We defined (i) **“long-lived”** as new spines that appeared and remained in place for each subsequent imaging session and (ii) **“short-lived”** as new spines that appeared but disappeared at subsequent imaging sessions. In some cases (< 10%), in which a spine appeared to be eliminated but reappeared in a subsequent session in the same location, that spine was considered short-lived. To this end, we first, counted all newly formed dendritic spines at days 13, 15, 18, and 25 (**Figure 3A**). In each imaging session, we observed a similar number of newly generated spines that was ∼3-4 fold greater in animals that received CHI (red curve) than in those that had received sham surgery (light blue curve). Remarkably, over the time-course of ISRIB treatment the number of newly formed spines normalized to control levels (dark blue curve) (**Figure 3B**). ISRIB treatment had no effect on the persistence of new spines in the control animals (**Figure 3B**).

**Figure 3.**
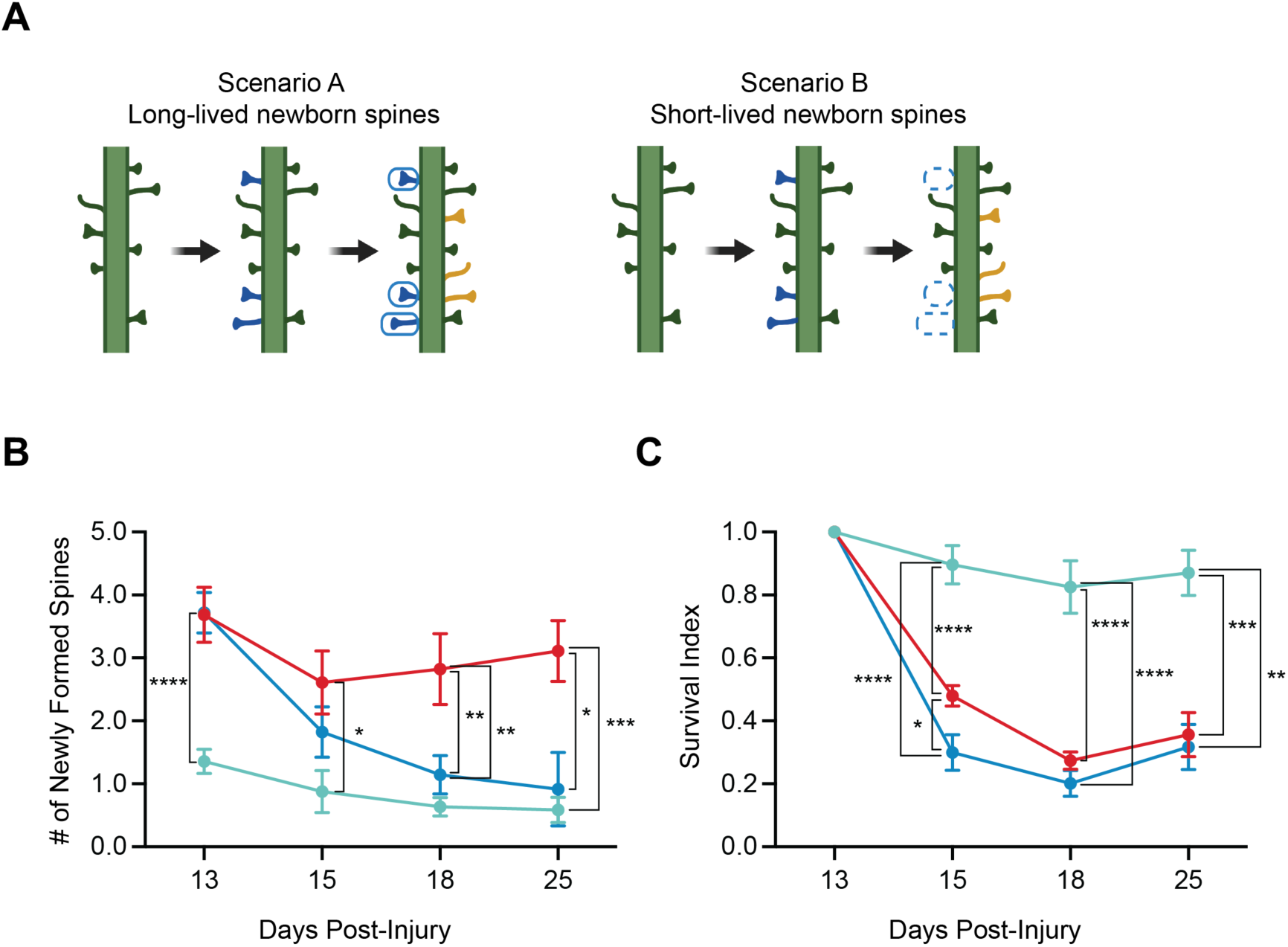
ISR inhibition normalizes the CHI-induced short-lived newborn spines. **(A)** A schematic illustrating long- vs. short-lived newborn spines. **(B)** Number of newly formed spines and **(C)** survival index in sham ± ISRIB (light blue line), CHI (red line), and CHI + ISRIB (dark blue line) mice (note that the CHI mice were spilt into the ± ISRIB groups) at day 13-25 post-injury (dpi). Sham ± ISRIB treatment showed no significant difference and were pooled. Statistics were calculated between the experimental groups and the pooled sham ± ISRIB treatment. ISRIB treatment had no effect on sham control mice (data not shown). **Statistics:** Analysis was done at each dpi by Unpaired T-test (B: 13 dpi) or Ordinary one-way ANOVA followed by multiple comparisons using Tukey-post hoc (B: 15-25 dpi, C: 15-25 dpi). p < 0.05 (*), p < 0.005 (**), p < 0.0005 (***), p < 0.00005 (****), p < 0.0001 (****), as indicated in the figure. All data are means ± SEM. Sham ± ISRIB n = 13; CHI n = 12; CHI + ISRIB n = 11.

Statistical analysis confirmed that CHI mice had significantly more newly formed spines than controls at each imaging session (p < 0.0001, p < 0.05, p < 0.005, p < 0.0005, respectively); conversely, ISRIB treatment of CHI animals significantly reduced newly formed spines, which became statistically indistinguishable from controls (p = 0.371, p = 0.440, p = 0.803) (**Supp Figure 5**). Thus, taken together, the data in **Figure 3** strongly suggest that most newly formed spines are short-lived.

To further test this conjecture, we determined the stability of each newly formed spine by notating whether it persisted in subsequent imaging sessions and calculating a **“survival index”**, in which a value closer to 1 denotes a larger fraction of long-lived spines and a value closer to 0 denotes a larger fraction of short-lived spines (*51*). New spines born between days 11 and 13, that is “new spines”, had a survival index of ∼0.9 in control animals when observed at day 15; this index was significantly smaller, ∼0.4, following CHI (p < 0.0005) (compare light blue and red curves, **Figure 3C**), indicating that the new spines in CHI animals added between these two imaging sessions were mostly short-lived. This difference was similar when indexed on days 18 and 25 post-injury (p < 0.0005, p < 0.005, respectively). Similarly, spines born between days 13 and 15 and indexed on days 18 and 25, showed a low survival index following CHI (**Supp Figure 6**). ISRIB treatment only affected the survival index at day 15 (curves red and dark blue; p < 0.05, p = 0.7495, p = 0.9611), underscoring the notion that spines newly formed after CHI are short-lived and quickly eliminated. ISRIB treatment normalized the number of CHI-induced spines toward control levels. The normalization effect lasted for at least one week after treatment was ended.

### ISR inhibition lastingly reverses working memory deficits measured after concussive injury

To assess functional consequences of the aberrant spine dynamics measured after CHI in the parietal cortex, we next carried out behavioral experiments blind to injury status and treatment to measure working memory, which is known to depend on parietal cortex function. To this end, we subjected mice to CHI as previously described and measured their performance in the novel object recognition task (NORT). NORT is commonly used to measure working memory in rodents, taking advantage of the mouse’s innate preference for novelty (*52, 53*). Briefly, on days 16 - 17 post-CHI, mice were individually habituated to the measurement arena for 10 minutes each day and were then exposed on day 18 post-CHI to two identical objects (“learning phase”). Mice were then removed from the arena for 5 minutes, during which time one of the now familiar objects was replaced with a novel object, and were then returned to the arena to explore and interact with the objects (“testing phase”) (blue arrowheads in **Figure 1**). Automated tracking software recorded the total interaction time with each object during the learning and testing phases and calculated a **“discrimination index”** (DI) for each phase (DI = (Time spent with novel – time spent with familiar) / total time). The DI from the testing phase was used as a metric of working memory function. A high DI score reveals discrimination between the novel and familiar object, and a low DI score indicates impaired discrimination, presumably as consequence of a defect in working memory.

In the testing phase, CHI mice had a significantly lower DI compared to controls, indicating that CHI induced deficits in working memory (p < 0.0005) (compare light blue and red dots; **Figure 4A**).

**Figure 4.**
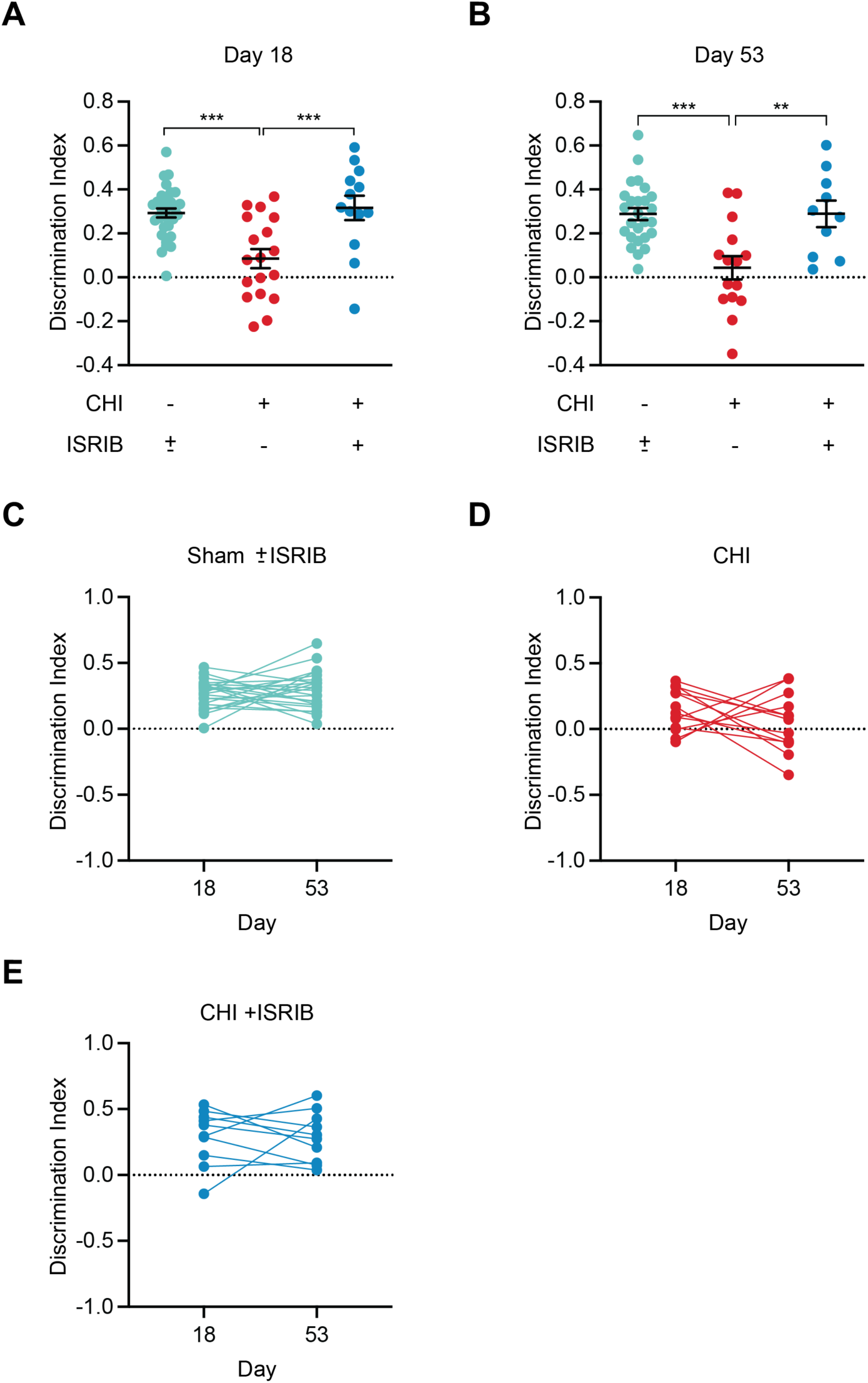
ISR inhibition lastingly reverses working memory deficits measured after concussive injury. Mice were treated as described before and subjected to cognitive testing on days 18 and 53 using NORT (see Methods) as depicted in Figure 1. The habituation phases of the NORT were performed over two days preceding days 18 and 53. **(A-B)** The discrimination index at day 18 (A) or day 53 (B) for sham ± ISRIB, CHI, and CHI + ISRIB mice. For both (A) and (B), sham ± ISRIB treatment showed no significant difference and were pooled. Statistics were calculated between the experimental groups and the pooled sham ± ISRIB treatment. ISRIB treatment had no effect on sham control mice (data not shown). **(C-E)** Connected dot plot comparing the discrimination index for individual animals at days 18 and 53 for sham ± ISRIB, CHI, and CHI + ISRIB mice. **Statistics:** Analysis was done by Ordinary one-way ANOVA followed by multiple comparisons using Tukey-post hoc (A-B) or Paired T-test (C-E). p < 0.005 (**), p < 0.0005 (***), as indicated in the figure. Data in (A) and (B) are means ± SEM. Sham ± ISRIB n = 31; CHI n = 18; CHI + ISRIB n = 13.

Using the same treatment regime described above for the spine measurements, we next tested whether ISRIB treatment would reverse the working memory deficits. Remarkably, CHI mice that received ISRIB treatment regained full working memory, with discrimination indistinguishable from that of the control group (p = 0.896) (compare light blue and dark blue dots; **Figure 4A**). ISRIB treatment did not affect the performance of the control animals (**Supp Figure 7, Figure 4C**).

We then repeated the NORT 35 days after the last ISRIB treatment in the same animal cohort but using a different novel object comparable in novelty (blue arrowhead on day 53 in **Figure 1**). We observed that CHI mice that had received ISRIB between days 14 and 17 post-CHI maintained full working memory at day 53 (p = 0.9911 n.s., comparing DI of day 18 with day 53) (**Figure 4B, E**). CHI mice that did not receive ISRIB remained impaired (p = 0.3097 n.s.) (**Figure 4B, D**), indicative of long lasting working memory deficits induced by CHI. The persistence of the ameliorative effects of ISRIB treatment on working memory deficits induced by CHI for weeks after administration parallels the persistent effects of ISRIB in restoring normal spine dynamics.

## Discussion

We demonstrate here that concussive injury induces aberrant spine dynamics in the parietal cortex that parallels impairments in working memory. Specifically, in each imaging session, we observed a significant increase in spine formation. Newly formed spines, however, were short-lived, as indicated by their short ∼2-day lifetime, suggesting that they do not engage in stable, functional synaptic connections. Remarkably, pharmacological inhibition of the ISR with ISRIB reversed the changes in spine dynamics and restored working memory. These restorative effects were maintained weeks after ISRIB treatment was terminated; that is, long after the drug was cleared from the organism. The strong correlation between the anatomical and behavioral findings, including their responses to ISRIB treatment, suggests that structural changes in dendritic spines may be causally linked to cognitive impairments after CHI.

Structural plasticity of dendritic spines is tightly coordinated with synaptic function and neuronal plasticity. For example, spine enlargement parallels long-term potentiation, whereas long-term depression is associated with spine shrinkage (*30, 54, 55*). Changes in dendritic spines can have marked effects on synaptic function, plasticity, and patterns of connectivity in neuronal circuits (*21, 29*). Notably, disruptions in dendritic spine dynamics, density, or morphology accompany many brain disorders, particularly those that involve deficits in cognitive function (*56–58*). Indeed, in an animal model of Huntington’s disease, spine formation increases but newly formed spines do not persist and do not become integrated with the local circuitry (*56*). Similarly, in mice overexpressing MECP2, a Rett Syndrome related gene, both spine formation and elimination are elevated, and new spines are more rapidly eliminated than in wild type mice (*57*).

Increasing evidence from animal studies demonstrates that TBI-induced functional deficits are closely related to damage to the synaptic connections of neurons, including dendritic spines (*59–63*). These studies show an acute reduction in dendritic spine density associated with impaired electrophysiological activity and cognitive function. One study using a pneumatic piston to deliver a single TBI in mice found an acute steady reduction in spine density of layer II/III pyramidal neurons within the parietal cortex (over 1 - 24 hours post-injury) that was followed by a slower increase in spine density (at 3 days post-injury) (*64*). Similarly, in another study analyzing the same region using a fluid percussion injury in rats, spine density declined acutely over 24 hours and then returned to control levels within a week post-injury (*65*). Thus, in each case, an initial acute reduction in spine density was followed by a slower rebounding phase that restored spine density to (or close to) pre-injury levels. However, these and other studies remained limited to focal brain injury models, fixed brain tissues, and single time-point analyses. In contrast to previous work, to our knowledge, this is the first study to use longitudinal imaging *in vivo* to reveal spine dynamics spanning an extended time window after concussive injury when persistent cognitive deficits are measured. Although the injury model, animal, mode of analysis, and time- points were different, we also observed the previously noted spontaneous rebound in dendritic spines in CHI. Our mode of analysis spanned an extended time window (spread over an 8 to 15 day period), allowing kinetic characterization of dendritic spine dynamics and density. It remains to be determined whether these observations extend to other brain cell types and regions.

Spine imaging was performed on anesthetized mice because of the required mechanical stability when imaging small structures at diffraction-limited resolution. For events that happen on a scale of days, this is a reasonable approach since the time under anesthesia is short (∼ 1 h) when compared to the time awake between imaging sessions. Thus, the changes observed largely occurred when the animal was awake. Due to the shallow tissue penetration afforded by the two-photon imaging modality, we restricted our analysis to apical dendrites in the superficial layers of the parietal cortex (encompassing the impacted region), where neurons that engage in working memory are positioned close to the brain’s surface.

Persistent deficits in memory and executive function after TBI have been observed in both humans and animal models (*48, 66–69*). Unlike moderate and severe TBI, that is, focal injuries leading to pronounced tissue damage visible by computed tomography (CT), mild TBI (concussive or repetitive rapid acceleration treatment) does not evoke detectable tissue lesions in the brain that can be detected by CT or magnetic resonance imaging (MRI). However, like severe TBI, mild TBI leads to long-term cognitive dysfunction. The CHI protocol used here resulted in working memory deficits that lasted at least two months after injury.

We and others previously showed that both focal and diffuse head injuries, various neurodegenerative diseases, and normal aging all induce phosphorylation of eIF2*α* in the brain and thereby activate the ISR (*8, 9, 70-75*). After TBI, temporary pharmacological ISR inhibition of the ISR with ISRIB completely rescued trauma-induced cognitive and behavioral deficits and neuronal correlates (*8, 9*). Our observation that ISRIB treatment not only rescued the working memory deficit but also rapidly and persistently reversed the aberrant changes in cortical spine dynamics suggests that the link between the ISR and memory function involves at least in part alteration in neuronal structure. In this view, the appearance of transient spines results from ISR activation, which in turn leads to the observed memory deficits because proper synaptic connections are not formed. It will be exciting to next explore whether these observations also extend to the hippocampus, the brain region in which long-term memory consolidation is thought to take place and where similarly remarkable curative effects of ISRIB have been documented.

## Materials and Methods

### Animals

All experiments were conducted in accordance with National Institutes of Health (NIH) Guide for the Care and Use of Laboratory Animals and approved by the Institutional Animal Care and Use Committee of the University of California, San Francisco. Male C57B6/J wild-type (WT) mice were received from Jackson Laboratories. Male Thy-1-YFP-H (in C57 background) were bred in house. Animals were 10-16 weeks of age at the time of surgeries. Animal shipments were received at least one week prior to start of experimentation to allow animals to habituate to the new surroundings. Mice were group-housed in environmentally controlled conditions with a reverse light cycle (12:12 h light: dark cycle at 21 ± 1 °C; ∼50% humidity) and provided food and water *ad libitum*. Behavioral analysis was performed during the dark cycle.

### Closed-head injury and Sham surgery

10-12 weeks old C57BL/6J or Thy-1-YFP-H (in C57 background) mice were randomly assigned to each CHI or sham surgery group. Animals were anesthetized and maintained at 2-2.5% isoflurane during CHI or sham surgery. Animals were secured to a stereotaxic frame with nontraumatic ear bars, and the head of the animal was supported with foam before injury. Contusion was induced using a 5-mm convex tip attached to an electromagnetic impactor (Leica) at the following coordinates: anteroposterior, −1.50 mm and mediolateral, 0 mm with respect to bregma. The contusion was produced with an impact depth of 1 mm from the surface of the skull with a velocity of 5.0 m/s sustained for 300 ms. No animals had a fractured skulls after injury. Sham animals were secured to a stereotaxic frame with nontraumatic ear bars and received the midline skin incision but no impact. After CHI or sham surgery, the scalp was sutured, and the animal was allowed to recover in an incubation chamber set to 37°C. All animals recovered from the surgical procedures as exhibited by normal behavior and weight maintenance monitored throughout the duration of the experiments.

### Drug Administration

ISRIB solution was made by dissolving 5 mg ISRIB (obtained from Peter Walter’s group) in 1 mL dimethyl sulfoxide (DMSO) (PanReac AppliChem, 191954.1611). The solution was gently heated in a 40 °C water bath and vortexed every 30 s until the solution became clear. Next 1 mL of Tween 80 (Sigma Aldrich, P8074) was added, solution was gently heated in a 40 °C water bath and vortexed every 30 s until the solution became clear. Next, 10 mL of polyethylene glycol 400 (PEG400) (PanReac AppliChem, 142436.1611), solution was added gently heated in a 40 °C water bath and vortexed every 30 s until the solution became clear. Finally, 36.5 mL of 5% dextrose (Hospira, RL-3040) was added. The solution was kept at room temperature throughout the experiment. Each solution was used for injections up to 7 d maximum. The vehicle solution consisted of the same chemical composition and concentration (DMSO, Tween 80, PEG400 and 5% dextrose). Stock ISRIB solution was at 0.1 mg/ml and injected at a dose of 2.5 mg/kg.

### Headplate and cranial window attachment surgery

10-12 weeks old Thy-1-YFP-H (in C57 background) mice underwent headplate and cranial window attachment surgery. Animals were anesthetized, maintained at 1.5% isoflurane, and placed over a protected thermal plate (37°C) during surgery. Animals were secured to a stereotaxic frame with nontraumatic ear bars. After anesthesia induction, dexamethasone (intraperitoneal), meloxicam (intraperitoneal), and lidocaine (subcutaneous) drugs were administered. The scalp was cleaned with ethanol 70%, disinfected with Betadine, and the skull was exposed with a midline scalp incision. A craniotomy was performed on the right parietal cortex with a high-speed drill equipped with a round bur. To avoid damaging the underlying cortex by friction-induced heat, a cool sterile solution was added to the skull periodically, and drilling was intermittent to permit heat dissipation. The excised skull was replaced by a 3 mm coverslip window (Harvard Apparatus, Round Cover Glass) carefully positioned on top of the brain and secured against the skull with tissue adhesive (3M Vetbond tissue adhesive). A custom-made headplate with a central opening was attached with dental cement (Parkell C&B Metbond Quick Adhesive Cement System). After the surgery, the animal was allowed to recover in an incubation chamber set to 37°C before returning to its home cage.

All animals recovered from the surgical procedures as exhibited by normal behavior and weight maintenance monitored throughout the duration of the experiments. Animals were allowed to recover before the start of imaging experiments.

### Transcranial two-photon imaging *in vivo*

Transcranial two-photon imaging was performed *in vivo* using a movable objective microscope manufactured by the Sutter Instrument Company. A mode-locked Ti:Sapphire laser (Chameleon Ultra 2, Coherent, Inc.) was tuned to 920 nm, and the laser power through the objective was adjusted within the range of 60 to 80 mW. Emission light was collected by a 40x water-immersion objective (NA0.8; IR2; Olympus), filtered by emission filters (525/70 and 610/75 nm; Chroma Technology) and measured by 2 independent photomultiplier tubes (Hamamatsu R3896). Scanning and image acquisition of apical dendrites in layer I/II of layer V pyramidal neurons expressing green fluorescent protein was controlled by ScanImage software (Vidrio Technologies). Z-stacks were collected with a step size of 1 µm in the *z* axis and the pixel size to 0.1513 µm. Each slice of the stack was obtained by averaging 25 frames. Mice were imaged for 4 or 5 sessions over an 8 or 15 day period. Care was taken with each imaging session to achieve similar fluorescence levels. Mice were held under isoflurane anesthesia (1.5% in oxygen) for one hour with head fixed during structural imaging. The body temperature of the mouse was kept at 37°C using a feedback-controlled electric heating pad.

### Spine dynamics analysis

For spine remodeling analysis, image stacks collected on day 13, 15, 18, and 25 were first aligned to their corresponding baseline imaging stack collected on day 11 using CMTK Registration Gui plugin to NIH ImageJ FIJI software (*76, 77*). Spine formation and elimination analysis was done semi-automated using the NIH FIJI software (ROI coordinates) to save the X, Y, Z coordinates of identified dendrites on traced dendrites from day 11 image stacks. The X, Y, Z coordinates were then applied to image stacks from subsequent imaging sessions to note the continued presence or disappearance of pre-existing spines or the presence of a new spine. Spine formation and elimination were graphed as compared to baseline imaging day 11.

40 – 57 dendrites ranging between 50-100 µm (+/- 2 µm) in length were analyzed per mouse and then averaged. A total length of at least 3500 µm of dendrites and 200-600 spines were analyzed per experimental group. To exclude the possibility that the measured changes analyzed were due to the choice of day 11 as the baseline for analysis, spine formation and elimination were analyzed in reverse by comparing days 11, 13, and 15 to day 18.

The number of new spines formed per imaging session was quantified by noting the number of new spines formed (first observed) at imaging day 13, 15, and 18. Multiple dendrites were analyzed per mouse and averaged.

To calculate the survival index, we determined the number of spines present at subsequent imaging sessions divided by the number of spines present when first observed. First, individual spines were classified as S_11_, S_13_, S_15_, or S_18_ depending on which imaging day they were first observed. Next, a binary system was used to denote presence (indicated by a 1) or not (indicated by a 0) across the imaging sessions. In this scale, a survival index closer to 1 denotes “long-lived” and a survival index closer to 0 denotes “short-lived.” For every dendrite, the survival index was calculated for each dendritic spine population (S_11_, S_13_, S_15_, or S_18_). The average survival index of each dendritic spine population was graphed per mouse.

### Spine density quantification

For spine density analysis (number of spines per µm of dendritic length), we determine the total number of spines present at baseline imaging day 11 divided by total number of spines present at each subsequent imaging session per dendrite. All protrusions greater than 0.55 µm from a dendrite were counted as spines regardless of morphology. The average spine density at imaging session was graphed per mouse.

### Tissue collection

Mice were lethally overdosed using a mixture of ketamine (10 mg/ml) and xylaxine (1 mg/ml). Once animals were completely anesthetized, blood was extracted by cardiac puncture and animals were perfused with 1X phosphate buffer solution, pH 7.4 (Gibco, Big Cabin OK, -70011-044) until the livers were clear (∼1–2min). For measurements of drug concentration following PBS perfusion, the whole brain was rapidly removed, a hemi-brain was dissected and weighed, and snap frozen on dry ice and stored at −80 °C until processing.

### Immunohistochemistry analysis and quantification

For immunohistochemistry analysis, following PBS perfusion, whole brains were fixed in ice-cold 4% paraformaldehyde, pH 7.5 (PFA, Sigma Aldrich, St. Louis, MO, 441244) for 4 h followed by sucrose (Fisher Science Education, Nazareth, PA, S25590A) protection (15% to 30%). Brains were embedded with 30% sucrose/Optimal Cutting Temperature Compound (Tissue Tek, Radnor, PA, 4583) mixture on dry ice and stored at -80 °C. Brains were sectioned into 20 μm slides using a Leica cryostat (Leica Microsystems, Wetzlar, Germany) and mounted on slides (ThermoFisher Scientific, South San Francisco, CA). Slides were brought to room temperature (20 °C) prior to use. Slides were washed in tris buffered saline tween solution for 10 minutes and twice with tris buffered saline (TBS) for 10 minutes each. All slides were blocked in Block Reagent (Perkin Elmer, Waltham, MA, FP1020) for 30 minutes in the dark. Slides were stained with primary antibodies specific for NeuN (Rabbit, Abcam, Burlingame, CA ab128886) overnight, washed three times in TBS, and stained for the secondary antibody, goat anti-rabbit Alexa-568 (Invitrogen, Carlsbad, CA, A- 11011). Tissues were fixed using ProLong Gold (Invitrogen, Carlsbad, CA, P36930) and a standard slide cover sealed with nail polish. 3-4 images separated by 60-140 µm in the cortex were averaged per animal. 15 µm z-stack images were acquired on a Zeiss Axio Imager Z1 fluorescence scope at 10x and 20x magnification. Quantitative analysis was performed using NIH FIJI analysis software (v1.52n). Neurons were determined using the neuronal marker (NeuN).

The neuronal numbers were determined by quantification of a neuronal nuclei and by the image- covering staining and expressed as total numbers or percentage of the total area.

### Novel object recognition task with 5-minute retention interval

For all behavioral assays the experimenters were blinded to injury and therapeutic intervention. Prior to behavioral analysis animals were inspected for gross motor impairments. Animals were inspected for whisker loss, limb immobility (included grip strength), and eye occlusions. If animals displayed any of these impairments, they were excluded. Behavioral assessment was recorded and scored using a video tracking and analysis setup (Ethovision XT 8.5, Noldus Information Technology).

The novel objective recognition task was modified from a previously described protocol [78]. The task took place during the dark cycle in a room with dim red light. Mice were individually habituated in a 30 cm × 30 cm × 30 cm (L×W×H) opaque plexiglass box (termed ‘arena’) for 10 minutes per day for 2 days preceding the test. On the third day, mice were exposed to two identical objects for 5 minutes (learning phase). The mice were then temporarily removed for 5 minutes to allow for one of the objects to be replaced by a novel object. Mice were then returned to the arena for 5 minutes to explore and interact with the objects (testing phase). In each experiment, the location of the novel object was randomly selected between the left and right side to avoid side-preference bias. Mouse behavior was recorded with an overhead video camera; the video files were exported to Ethovision XT 8.5, Noldus Information Technology for analysis. Data are presented as a discrimination index (DI), calculated using formula DI = ((Time spent with novel – time spent with familiar)/total time).

Animals were injected (intraperitoneal) with either vehicle or ISRIB (2.5 mg/kg) starting two days before the first training day (days 14 and 15 post-injury) (**Figure 1**) and after each of the habituation days (days 16 and 17 post-injury) for a total of four doses. No injections were given when working memory was tested on day 18 post-injury or thereafter. On day 53 (that is, 35 days post the last ISRIB/vehicle treatment), we measured working memory as described above but used a different novel object comparable in novelty.

### Statistical Analysis

All data were analyzed with GraphPad Prism 9 statistical software. Unpaired T-test and One-way analysis of variance (ANOVA) (individual statistical tool and post-hoc analysis denoted in Figure Legends). The p values of < 0.05 were considered as significant. Individual animal scores represented by dots, lines depict Data mean and SEM. Group outliers were determined (GraphPad Software Outlier Test-Grubb’s test) and excluded from analysis. At most a single animal was excluded from each experimental cohort.

## Acknowledgments

This work was supported by National Institutes of Health grants R01AG056770 (SR), R01EY002874 and R01MH122478 (MPS), National Science Foundation Grant 1822650 (MPS), the generous support of the Roger’s family to SR and PW, the Weill Innovation Award to SR and PW. PW is an Investigator of the Howard Hughes Medical Institute. ESF was supported by the National Institute of General Medical Sciences (NIGMS) Initiative for Maximizing Student Development (IMSD) (R25GM056847) and the National Science Foundation (NSF) Graduate Research Fellowship Program (GRFP). Figures 1, 2A, and 3A were created with Biorender.com.

---

**Supplemental Figure 1.**
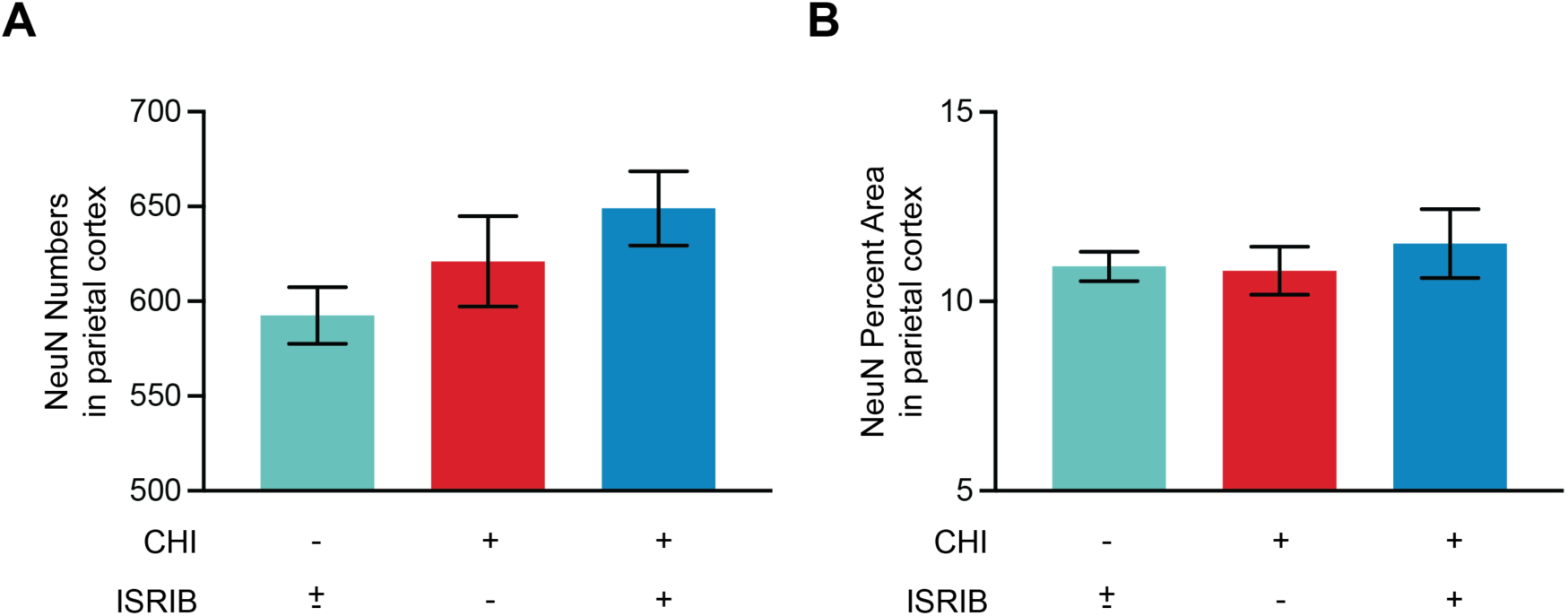
No changes in neuronal numbers or percent area in the parietal cortex following closed-head injury or ISRIB treatment. **(A-B)** Neuronal cell bodies were quantified in the parietal cortex from adult male WT mice. Neuronal nuclei protein NeuN was used to count neuronal cell bodies. No significant differences in NeuN numbers or percent area were found at 10x and 20x magnification. **Statistics:** Analysis was done by Ordinary one-way ANOVA followed by multiple comparisons using Tukey-post hoc (A-C). p < 0.0005 (****), as indicated in the figure. All data are means ± SEM. Sham ± ISRIB n = 5; CHI n = 4; CHI + ISRIB n = 4.

**Supplemental Figure 2.**
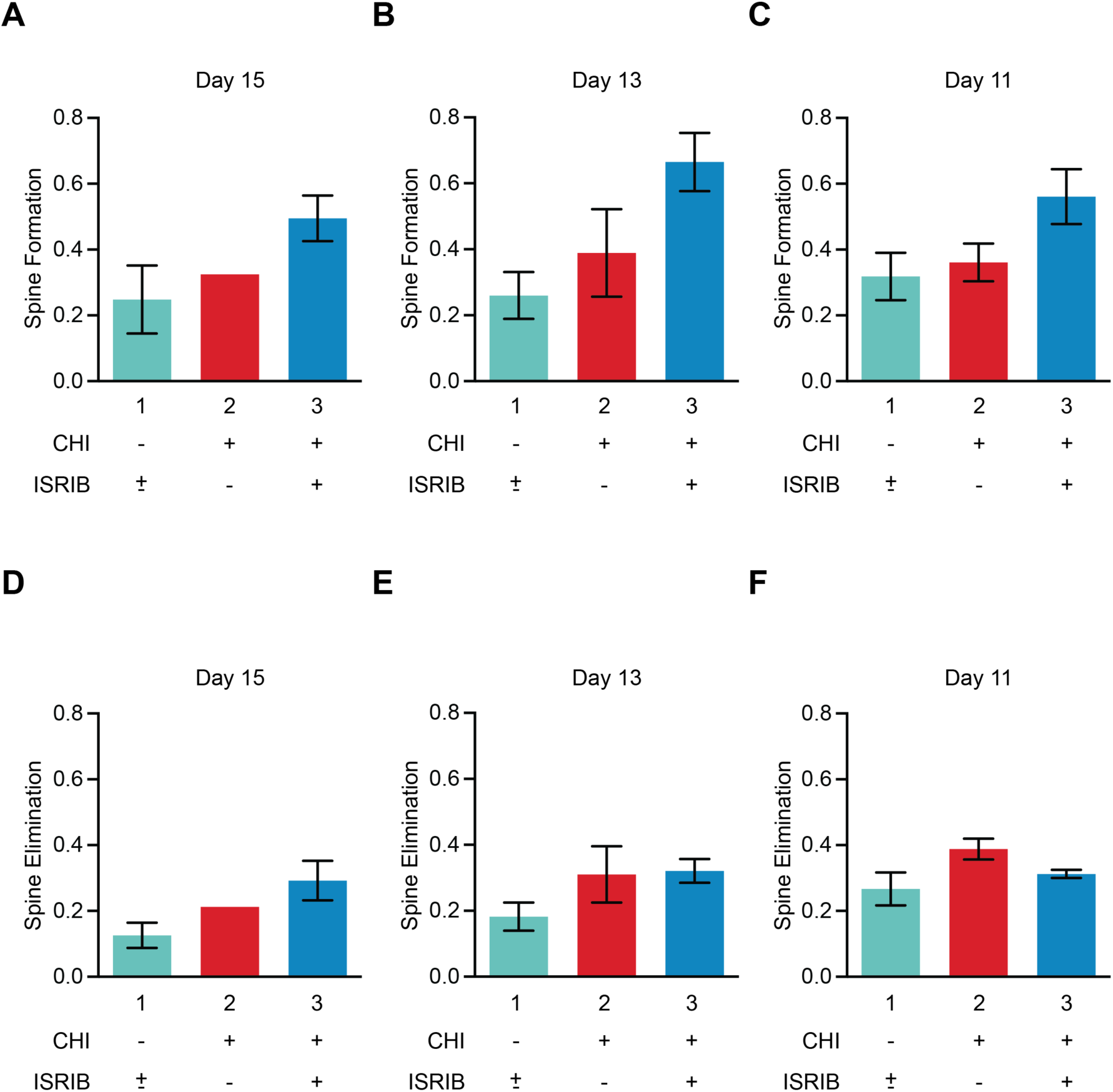
Reverse analysis using day 18 as the baseline imaging day. Reverse-time analysis of **(A-C)** spine formation and **(D-F)** elimination using day 18 as the baseline imaging day. All data are means ± SEM. Sham ± ISRIB n = 13; CHI n = 12; CHI + ISRIB n = 11.

**Supplemental Figure 3.**
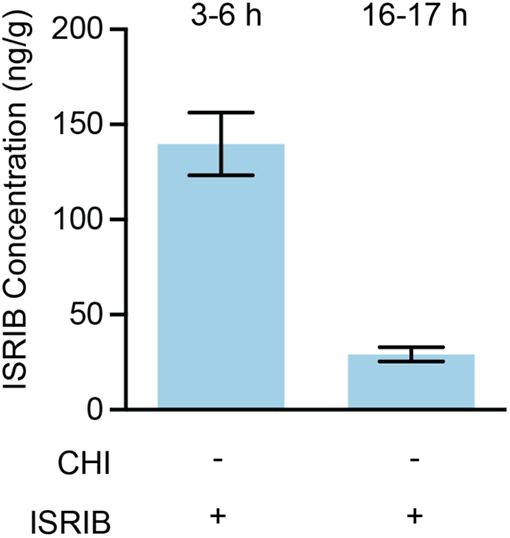
ISRIB concentration in the brain. ISRIB concentration (ng/g) in the brain of control WT mice at 3-6h or 16-17h post last injection. All data are means ± SEM. Sham ± ISRIB n = 12, 6, respectively.

**Supplemental Figure 4.**
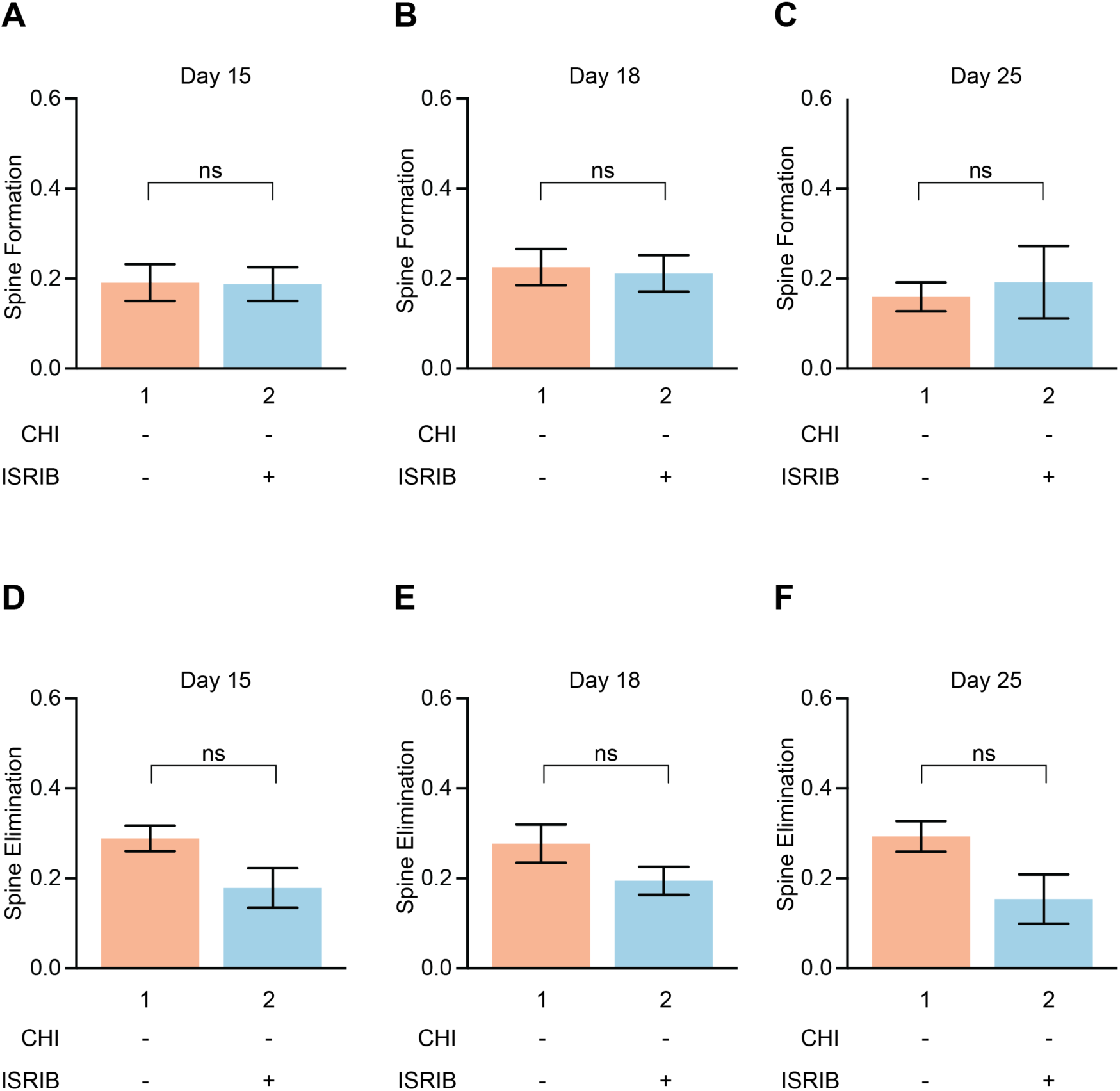
ISRIB treatment does not affect spine formation or elimination of sham control mice. **(A-C)** Spine formation and **(D-F)** spine elimination at day 15, 18, and 25 dpi for Sham – ISRIB (Bar 1) and Sham + ISRIB (Bar 2). ISRIB treatment had no effect on sham control mice. The number of animals in each group was n = 6 and 7, respectively. **Statistics:** Analysis was done by Unpaired T-test (A-F). All data are means ± SEM. Sham ± ISRIB n = 13.

**Supplemental Figure 5.**
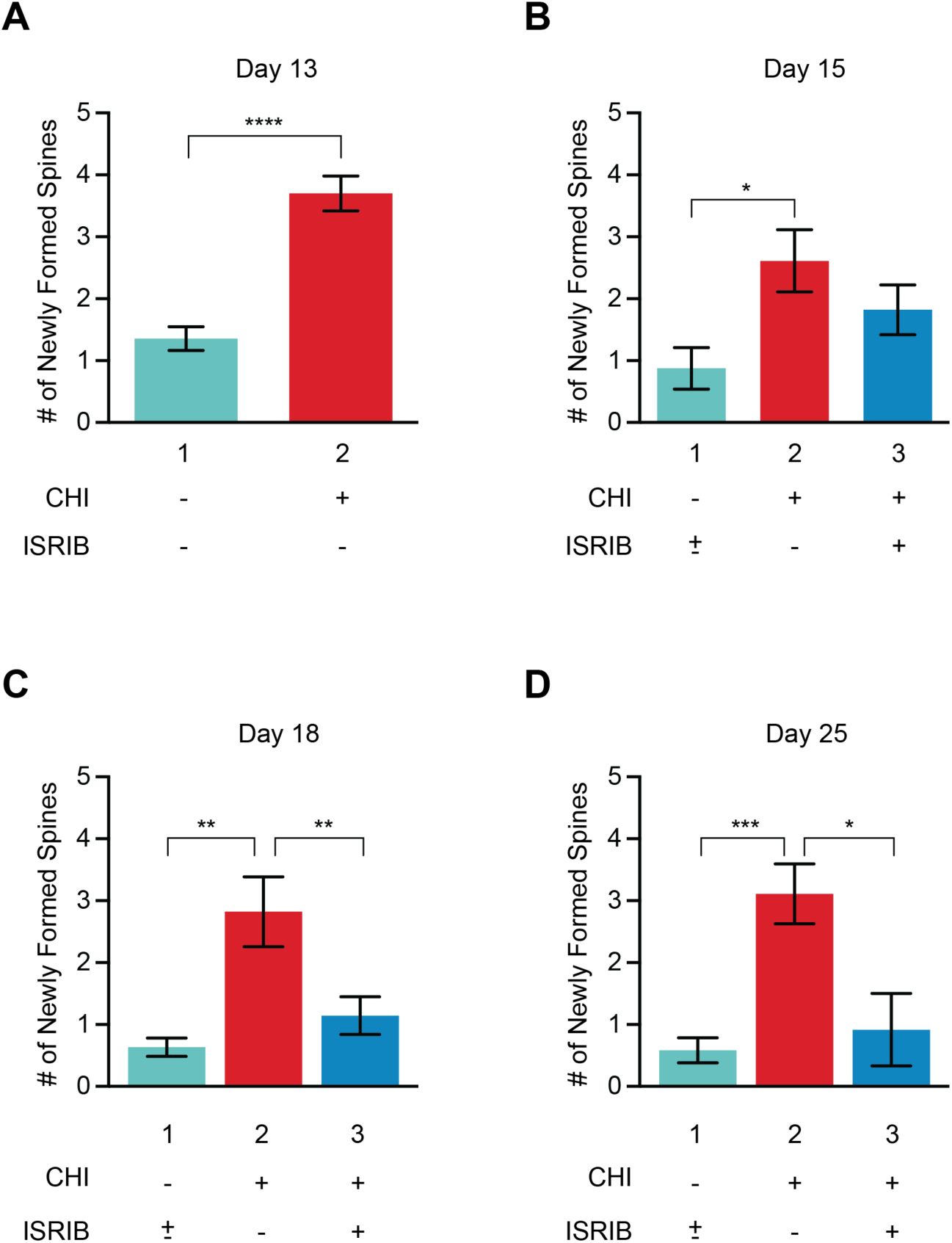
Number of newly formed spines at day 13-25 post-injury. **(A-D)** Number of newly formed spines in Sham ± ISRIB (Bar 1), CHI (Bar 2), and CHI + ISRIB (Bar 3) mice at day 13-25 post-injury (dpi). **Statistics:** Analysis was done by Unpaired T-test (A) or Ordinary one-way ANOVA followed by multiple comparisons using Tukey-post hoc (B-D). p < 0.05 (*), p < 0.005 (**), p < 0.0005 (***), p < 0.00005 (****), as indicated in the figure. All data are means ± SEM. Sham ± ISRIB n = 13; CHI n = 12; CHI + ISRIB n = 11.

**Supplemental Figure 6.**
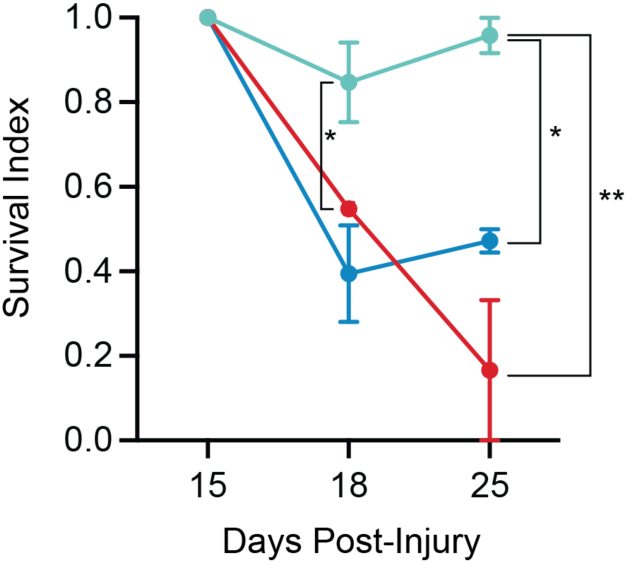
Survival index of spines born between days 13 and 15 and indexed on days 18 and 25 at day 15-25 post-injury. Survival index in Sham ± ISRIB (light blue line), CHI (red line), and CHI + ISRIB (dark blue line) mice at day 15-25 post-injury demonstrate same trends as seen in Figure 3C. **Statistics:** Analysis was done at each dpi by Ordinary one-way ANOVA followed by multiple comparisons using Tukey-post hoc. p < 0.05 (*), p < 0.005 (**), as indicated in the figure. All data are means ± SEM. Sham ± ISRIB n = 13; CHI n = 12; CHI + ISRIB n = 11.

**Supplemental Figure 7.**
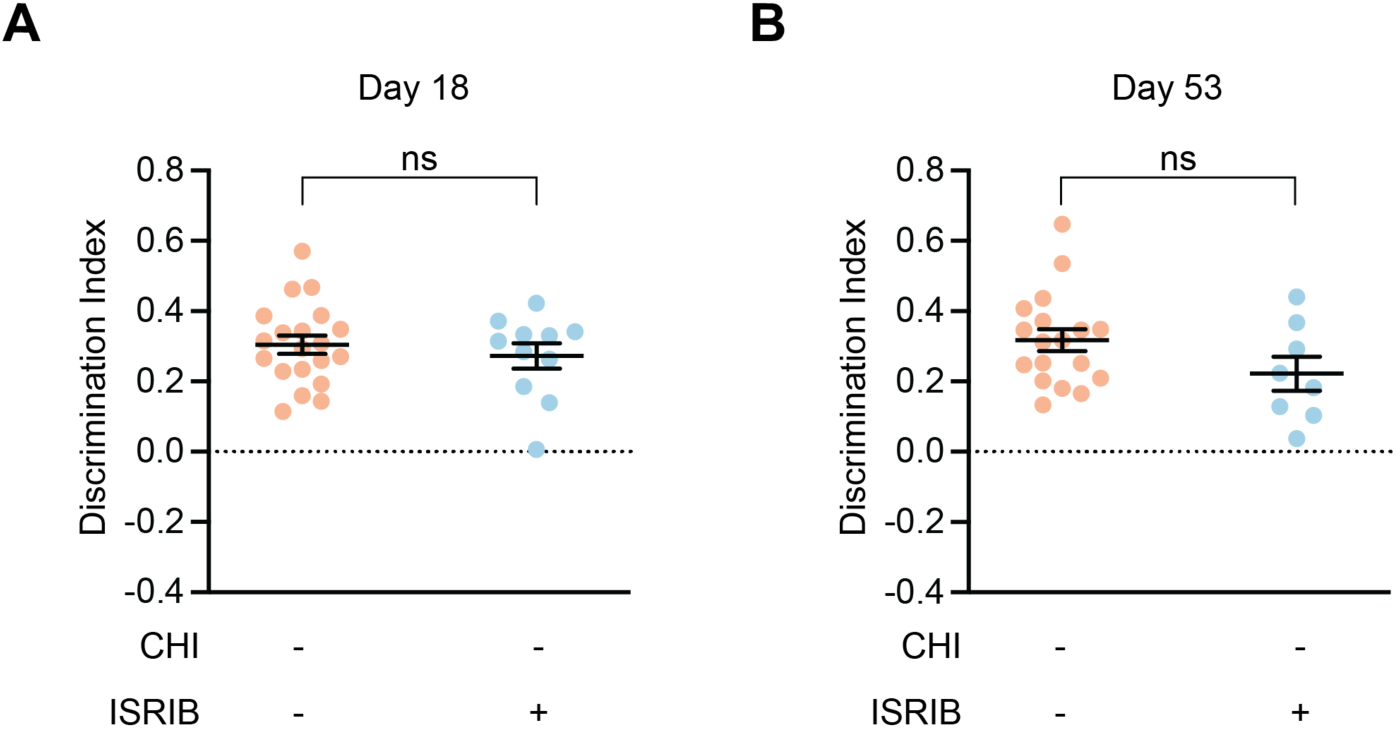
ISRIB treatment does not affect sham control mice performance on novel object recognition task. **(A-B)** The discrimination index at day 18 (A) or day 53 (B) for Sham ± ISRIB. For both (A) and (B), Sham ± ISRIB treatment showed no significant differences. **Statistics:** Analysis was done by Unpaired T-test (A-B). Data in (A) and (B) are means ± SEM. Sham ± ISRIB n = 31.

